# SARS-CoV-2 infects carotid arteries: implications for vascular disease and organ injury in COVID-19

**DOI:** 10.1101/2020.10.10.334458

**Authors:** Susanne Pfefferle, Thomas Günther, Victor G. Puelles, Fabian Heinrich, Dominik Nörz, Manja Czech-Sioli, Alexander Carsten, Susanne Krasemann, Milagros N. Wong, Lisa Oestereich, Tim Magnus, Lena Allweiss, Carolin Edler, Ann Sophie Schröder, Maura Dandri, Tobias B. Huber, Markus Glatzel, Klaus Püschel, Adam Grundhoff, Marc Lütgehetmann, Martin Aepfelbacher, Nicole Fischer

## Abstract

Stroke and central nervous system dysfunction are cardinal symptoms in critically ill corona virus disease 19 (COVID-19) patients. In an autopsy series of 32 COVID-19 patients, we investigated whether carotid arteries were infected with SARS-CoV-2 by employing genomic, virologic, histochemical and transcriptomic analyses. We show that SARS-CoV-2 productively infects and modulates vascular responses in carotid arteries. This finding has far reaching implications for the understanding and clinical treatment of COVID-19.

## Main

In December 2019, a cluster of pneumonia caused by a novel coronavirus (SARS-CoV-2) was reported in Wuhan, China. Corona virus disease 19 (COVID-19) has since spread worldwide affecting more than 7,750,000 people and causing more than 430,000 deaths by June 14, 2020. COVID-19 is characterized by respiratory symptoms with a significant number of patients developing acute respiratory distress syndrome (ARDS) and multi-organ failure, eventually leading to death, especially in older patients with multiple co-morbidities ^1–4^. In critically ill patients, who often present with viremia, the virus has been detected in multiple organs ^2^. A large autopsy series revealed that pulmonary artery embolism and/or deep vein thrombosis were present in about 40% of cases, indicating that COVID-19 is strongly associated with coagulopathy ^3^. Notably, large vessel stroke including one proven case of carotid artery thrombosis was reported in 5 younger COVID-19 patients ^5^. In 58 COVID-19 patients with ARDS, neurologic symptoms associated with brain perfusion abnormalities and/or stroke were highly prevalent ^6,7^. Endotheliitis as the potential cause for the COVID-19 induced vascular dysfunction and coagulopathy was inferred by histopathology analysis of various postmortem tissues from a limited number of patients ^8,9^.

Using real time PCR, we quantified SARS-CoV-2 RNA in Arteria carotis communis (A. carotis, carotid artery) samples and matching lung- and throat samples from 32 COVID-19 patients (basic clinical data see online methods). All patients showed high viral loads in the lung, consistent with viral pneumonia ^3^. In 26 (81.25%) of the A. carotis samples viral RNA was detected, with mean viral loads that were comparable to those observed in throat samples (Fig. 1a). Histopathologic findings in the carotid arteries were age-appropriate with a moderate degree of inflammation and no obvious signs of vasculitis (supplemental Table S1). On the basis of virus positivity in A. carotis and the availability of additional tissues and organs, we attempted live/infectious virus isolation from A. carotis and 10 matching organ samples (Fig, 1b). We included 7 patients in this analysis (Fig. 1a, red circles), out of whom 5 had ante-mortem viremia, one had no viremia and one had an unknown viremia status. The clinical histories of the patients are summarized in supplementary Table S2. Virus isolation most frequently (up to 75%) succeeded in A. carotis and Vena saphena as well as in lung, throat and jejunum, with the latter three tissues being known sites of virus replication ^10,11^. Viral genome analysis revealed identical viral sequences in A. carotis and lung samples from the same patients, whereas a number of single nucleotide variations differentiated sequences from different patients. All SARS-CoV-2 sequences belong to the most prevalent genotype in Europe (supplemental Fig S1). Isolation of infectious virus from organs required ante-mortem viremia and isolation rates correlated well with viral loads in the respective organs (Fig. 1b). Virus isolation from liver, myocardium, spleen and frontal cortex of the brain was unsuccessful. Relative abundance of subgenomic (sg) SARS-CoV-2 RNAs (which are generated during virus replication) were determined for 5 of the 7 carotid arteries (arteries #6 and #7 could not be sequenced due to low RNA quality) and were found to be, on average, comparable or even higher than those observed in lung samples (Fig. 1c, supplementary Table S3). Together these data indicate that SARS-CoV-2 infects and replicates in carotid arteries.

**Figure 1:**
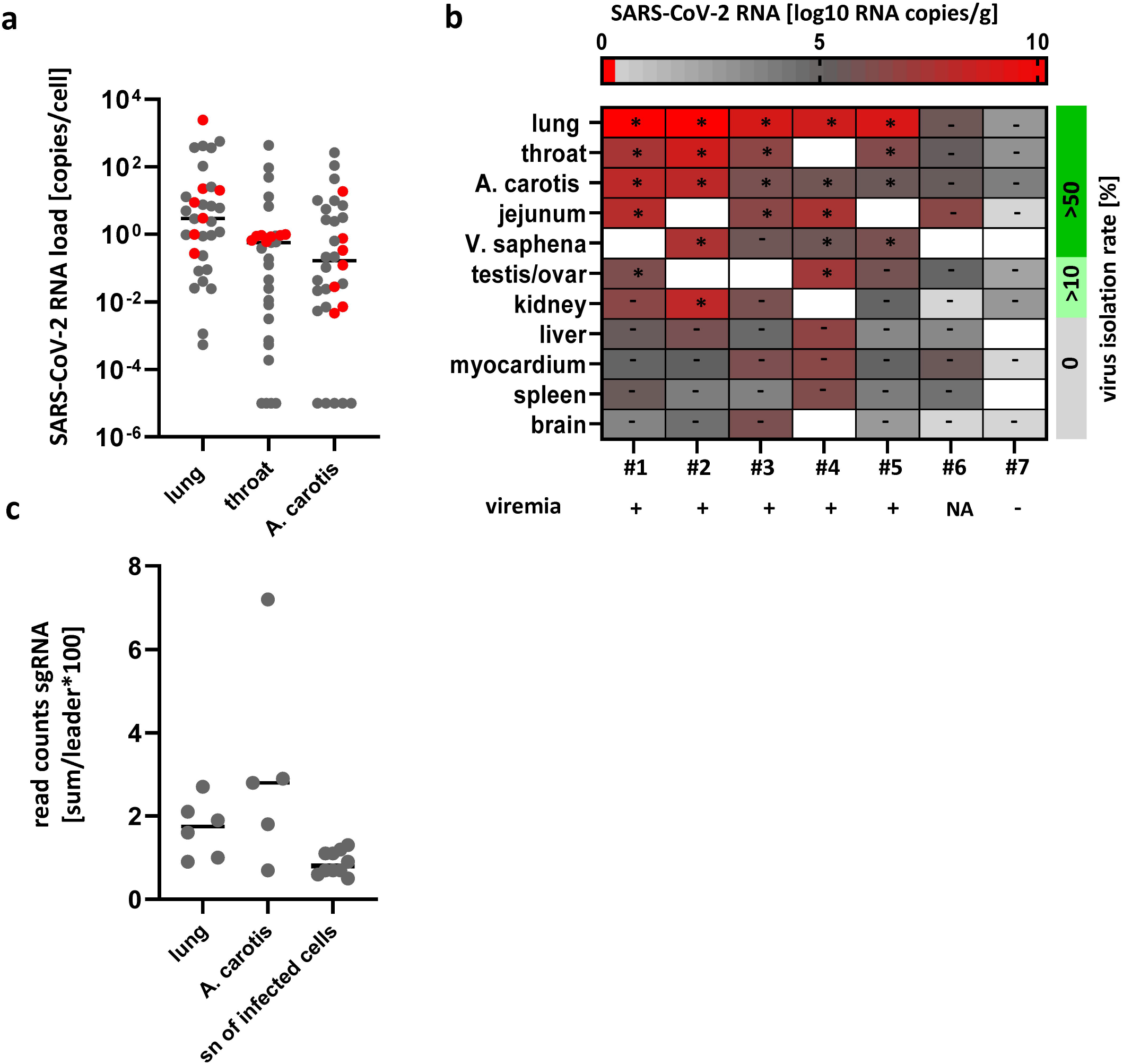
a) SARS-CoV-2 RNA concentrations in 32 post mortem lung, throat and A. carotis samples. Red dots indicate values of the 7 patients further investigated in b. (b) Virus isolation rates (* indicates successful isolation) and corresponding virus concentrations in indicated organs and ante mortem viremia in 7 patients (#1 - #7); nd: not dertermined. (c) Subgenomic (sg) SARS-CoV-2 RNA in 6 lung and 5 A. carotis samples and supernatants (sn) of SARS-CoV-2 infected Vero cells as negative control as determined by amplicon sequencing.

To investigate which cells in the carotid artery wall were infected with the virus, we performed SARS-CoV-2 plus-strand RNA in situ hybridization as well as anti-spike immunofluorescence and -immunohistochemistry in tissue sections. These staining techniques conclusively detected viral RNA and protein in the endothelial layer of the artery wall (Fig. 2a-d, supplemental Fig. S2).

**Figure 2:**
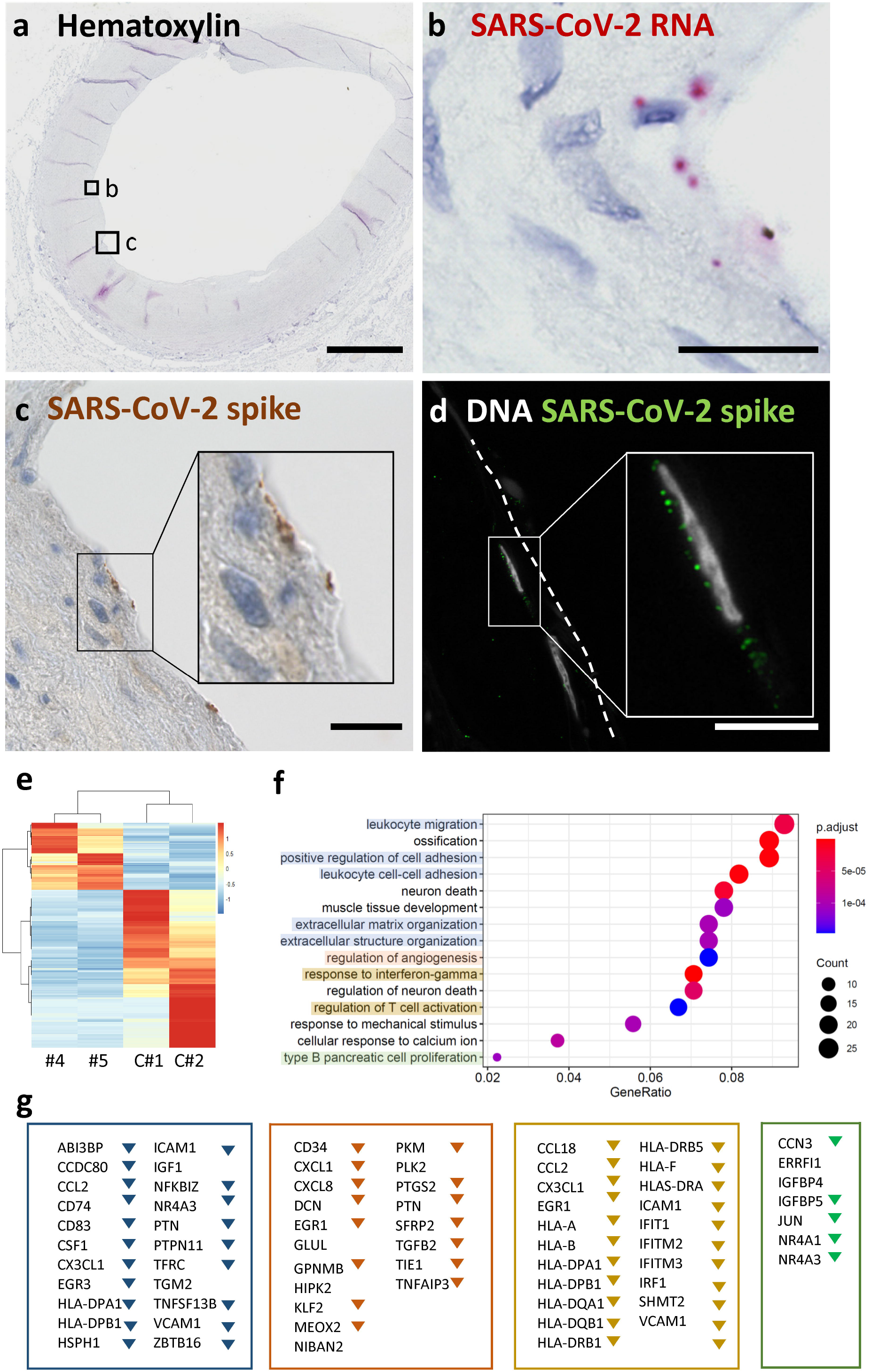
(a) Overview shows a cross section of an A. carotis counterstained with Hematoxylin and subjected to in situ SARS-CoV-2 RNA hybridization. (b) Close up of boxed region B in (a). (c) Close up of boxed region C, which depicts immunohistochemical staining of SARS-CoV-2 spike protein in a section consecutive to (a). (d) Immunofluorescence staining of nuclei and spike protein in endothelial cells seen in a section consecutive to the section shown in (a). Scale bars represent 300 μm (a) and 20 μm (b, c, d). (e) Transcriptional responses of A. carotis tissue from patients 4 and 5 and two patients, who died unrelated to COVID-19. Heat map of differentially expressed genes, DEGs (p-adjusted <0.1) with a log2FC >2. Down regulated genes are represented in gray to blue while up regulated genes range from yellow to red. Gene ontology analyses were performed using significantly regulated genes against the GO dataset for biological processes. The color of the dots represents the false discovery rate (FDR) value for each enriched GO term, and size represents the percentage of genes enriched in the total gene set. (f) Gene lists represent genes contributing to the GO terms extracellular matrix, angiogenesis, interferon response and proliferation. Down regulated genes are labelled with triangles.

We performed transcriptome analysis in A. carotis tissues from patients 4 and 5 (selected on basis of RNA quality) and two non-COVID-19 control patients in order to characterize the transcriptional response of SARS-CoV-2 infected arteries. Interestingly, type I and type II interferon pathways were down regulated in the SARS-CoV-2 infected arteries, which is in line with recent work demonstrating lowered interferon-related pathways and innate antiviral responses in SARS-CoV-2 infected cell lines and animal models ^12^. Of note, pathways and genes indicative of an inflamed endothelium (e.g. VCAM and NFκB) and an angiogenic vessel response (e.g. TIE1 and EGR1) were also strongly down regulated in the arteries of the deceased COVID-19 patients, suggesting that SARS-CoV-2 infection induces a strong anti-inflammatory and anti-proliferative state in blood vessels. Although these results comply with recent reports of a strong immunosuppressive effect of SARS-CoV-2 infection ^12^, they need to be interpreted with caution because patients 4 and 5 suffered from acute myeloid leukemia and myelofibrosis, respectively. These conditions are not generally associated with decreased inflammatory responses, but it is nevertheless possible that disease-specific immunodeficiency may have contributed to the observed phenotypes.

We propose that after entering the bloodstream, SARS-CoV-2 infects and replicates in vascular endothelial cells, where it modulates vascular responses and further migrates into organs. Notably, an increase in Kawasaki disease, a medium vessel vasculitis in children, has been recently linked to COVID-19 ^13,14^. How exactly infection of carotid arteries or other blood vessels contribute to the clinical symptoms, organ infection and final outcome of COVID-19 needs to be investigated in future studies.

## Acknowledgement

We thank D. Indenbirken and K. Hartmann for excellent technical support. We are grateful to all staff members of the institutes contributing to the study.

## Author’s contribution

S.P., T.G., V.G.P., D.N., M.C, M.W., F.H., K.P, S.K., L.O., L.A., A.F., M.L. performed experiments. S.P., T.G., K.P, M.G., M.D., T.B.H., N.F., A.G., T. M., M.A., M.L., L.O., L.A., A.F., M.L. contributed to the study design and data analysis. S.P., T.G., N.F., M.A., M. L. wrote the manuscript. T.G., K.P., T.M., M.G. and M.A interpreted clinical data. All authors approved the final version of the manuscript.

