## Supplementary Material for "SARS-CoV-2 infects carotid arteries: implications for vascular disease and organ injury in COVID-19"

**Supplementary online material**

- **Supplementary methods**
- **Supplementary Tables S1-S3**

Supplementary Table S1: Histopathological assessment of A.carotis communis samples.

Supplementary Table S2: Summary of preexisting comorbidities and postmortem findings of 7 patients included for detailed analysis

Supplementary Table S3: Subgenomic RNA detection by targeted amplicon sequencing.

- **Supplementary Figures S1-S3**

Supplementary Figure S1: Clustering of viral variants of SARS-CoV-2 sequences recovered from tissue sections of A. carotis and lung tissue.

Supplementary Figure S2: Control SARS-CoV-2 spike protein staining on tissue sections and infected Vero cells.

Supplementary Figure S3: Summary of gene expression analysis. Gene ontology and gene set enrichment analysis, GSEA.

**Supplementary methods:**

**Ethics:** The study was approved by the Ethics Committee of the Hamburg Chamber of Physicians (PV7311) and the local clinical institutional review board complying with the Declaration of Helsinki. Autopsies were mandated by the German federal state of Hamburg according to the German Infection Protection Act for all patients with PCR confirmed COVID-19.

**Patients and Autopsies:** 32 Patients dying of COVID-19 (PCR confirmed) and two COVID-19 negative controls (non-infection related cause of death) were included in the study. Clinical information including ante mortem diagnostic findings were obtained from patient’s medical records. All autopsies were performed under personal protection measures at the Institute of Legal Medicine, University Medical Center Hamburg-Eppendorf as recently described [1, 2]. The median age was 79 years (IQR 14.63), with 38% being female. The majority of cases (81%) represent hospitalized patients. Comorbidities within the cardiovascular, the respiratory, and the genito-urinary system were found in 90.6%, 65.6% and 59.4% of the cases, including arterial hypertension (88%), diabetes type 2 (34%), obesity (50%) and pre-existing chronic kidney failure (47%). 10 patients were immunocompromised due to pre-existing medical conditions or medication.

SARS-CoV-2 related pneumonia (as recently described in more detail [2]) was confirmed in 94% of the cases (n=30), in 69% being the alleged cause of death. 44% of patients suffered from deep vein thrombosis and 31% from pulmonary arterial embolism which is in line with previous published case series [1, 2]. 25% of the patients died of fulminant pulmonary embolism. Other causes of death were bronchitis and tracheitis (1/32) and infarction of the middle cerebral artery (1/32).

**Cell culture and virus isolation:** Vero cells (ATCC® CRL-1586) were cultivated in DMEM containing 10 % FCS, 1 % Penicillin/ Streptomycin, 1 % L-Glutamine, (200 mM), 1 % Sodium pyruvate, 1 % non-essential amino acids (all Gibco/ Thermo Fisher, Waltham, USA) under standard conditions. For virus isolation, cells were seeded into 24-well plates (TPP, Switzerland) at 80-90 % confluence. For infection, 250 µl of the homogenized tissue solution was added to each well and incubated for 1h at 37°C to allow for virus adsorption. Subsequently, cells were washed once with phosphate buffered saline (PBS) and 1 ml of fresh cell-culture medium was added. Cells were monitored daily for cytopathic effect. After 48-72 hours cell culture supernatant was harvested, filtered and stored at -80°C. Virus growth was confirmed by quantitative RT-PCR [3] and quantification by performing plaque-assays.

**Plaque assay:** Vero cells were seeded to confluence in 24-well plates. 200 µl each of a serial ten-fold virus dilution was used for infection. After adsorption, cells were overlaid with DMEM containing 1 % methylcellulose and incubated for 5 days. Plates were soaked for 30 min in a 4 % formaldehyde solution to allow for virus inactivation and cell fixation. Viral plaques were counted using crystal violet staining.

**Molecular diagnostic:** For detection of SARS-CoV-2 in tissue samples, nucleic acid extraction was performed automatically as described previously [4] by using either a Magna Pure Instrument (Roche, USA) or a QIASymphony instrument (Qiagen, Thermo Fisher, USA) and 200 µl of the homogenized tissue sample. A SARS-CoV-2 RT-PCR was performed on a Light Cycler 480 II instrument (Roche, USA) as described previously [5], using the 1 step RNA control kit (Roche, USA). As a marker for nucleic acid extraction quality in tissue samples, human ß-Globin gene was quantified by PCR (Life technologies, Thermo Fischer, USA). Detection of SARS-CoV-2 RNA in cell culture supernatant was performed as described previously [3].

**Histopathologic examination and immunofluorescence:** Formalin fixed paraffin embedded tissue (FFPE) samples were processed using a Ventana benchmark XT autostainer via standard procedure to slides stained with hematoxylin–eosin. The degree of arteriosclerosis, plaque formation and vessel associated lymphocytic infiltration was assessed using a semi quantitative approach.

Antibody for SARS-CoV-2 staining on FFPE tissue sections was validated on SARS-CoV-2 infected (HH-1 isolate [6]) and non-infected Vero cells that were fixed in formalin and embedded into paraffin. Anti-SARS-CoV-2 spike antibody (GeneTex; #GTX632604; clone 1A9) was used at 1:300 dilution and the abundance of infected cells were determined by experienced morphologists. Images were taken with a digital Leica DMD108 microscope. Immunofluorescence was performed using the anti-SARS-CoV-2 spike antibody described earlier at a 1:400 dilution as previously described [4].

**RNA in situ hybridization (RNAScope):** SARS-CoV-2 RNA was detected in formalin-fixed paraffin-embedded sections with the RNAScope 2.5 HD Duplex Assay (Advanced Cell Diagnostics, Newark, California, USA) as previously described [4]. The assay was performed according to the manufacturer´s instructions using standard pretreatment conditions. The probe V-nCoV2019-S (catalog number 848561-C2) specifically detected SARS-CoV-2 genomic RNA. Hematoxylin staining was performed to visualize nuclei. Stained sections were analyzed by fluorescence microscopy (Biorevo BZ-9000, Keyence, Osaka, Japan).

**SARS-CoV-2 amplicon sequencing:** Sample preparation for SARS-CoV-2 amplicon sequencing was based on the nCoV-2019 sequencing protocol from the Artic network project (artic.network/ncov-2019, dx.doi.org/10.17504/protocols.io.bbmuik6w) with the following modifications. Reverse transcription was performed in the presence of 2.4 µM random primer mix (mixture of hexamers and anchored-dT primer (dT23VN)) After RNase H digestion multiplex PCR was performed according to the Artic nCoV-2019 protocol using the primer scheme version 3 with 196 primers generating 400bp amplicons. Adaptor ligation for Illumina compatible multiplex sequencing was achieved by using the NEBNext DNA Ultra II Library Prep Kit for Illumina and the NEBNext Multiplex Oligos (96 Unique Dual Index Primer Pairs; New England Biolabs). All samples were multiplexed sequenced on Illumina MiSeq using 500cycle MiSeq v2 reagent kits (Illumina).

**Bioinformatic analysis of amplicon sequencing:** Illumina paired end amplicon sequencing reads were first merged using pear [7]. Merged reads were aligned to NC_045512.2 using minimap2 [8] with default settings for short read alignment. Merged reads were filtered by amplicon length and the presence of correct amplicon primer sequences at both ends. Variants were called using freebayes Bayesian haplotype caller v1.3.1 [9] with ploidy and haplotype independent detection parameters to generate frequency-based calls for all variants passing input thresholds (-K -F 0.01). Input thresholds were set to at least 10 variant supporting reads with a minimum base quality of 30 (-C10 -q30). Only high confidence variants present in > 33% of reads within at least one individual sample were included and annotated using ANNOVAR [10]. Minor frequency variants resulting in frame shift, stopgain or startloss were excluded. To detect subgenomic mRNAs we counted the reads containing 10 upstream bases and the 6 core element bases of the transcription-regulating sequence (TRS-L) fused to the 10 bases following the TRSs of the subgenomic mRNA bodies (TRS-B) of the individual subgenomic mRNAs.

**Transcriptome sequencing and bioinformatic analyses:** Extracted RNA (see above) was subjected to DNAse I digestion (Thermo Fisher Scientific) followed by RNeasy MinElute column clean up (Qiagen). Prior to Illumina compatible sequencing library generation, the RNA integrity was analyzed with the RNA 6000 Pico Chip on an Agilent 2100 Bioanalyzer (Agilent Technologies). rRNA was depleted (RiboCop rRNA Depletion Kit V1.2, Lexogen) and RNA-Seq libraries including unique molecular identifiers (UMIs) were generated using the CORALL Total RNA-Seq Kit (Lexogen) as per the manufacturer´s recommendations for degraded samples. Concentrations of all samples were measured with a Qubit 2.0 Fluorometer (Thermo Fisher Scientific) and fragment lengths distribution of the final libraries was analyzed with the DNA High Sensitivity Chip on an Agilent 2100 Bioanalyzer (Agilent Technologies). All samples were normalized to 2nM and pooled equimolar. The library pool was sequenced on the NextSeq500 (Illumina) with 1x75bp.

De-multiplexed libraries were aligned to the human reference genome hg38 using STAR v2.6.1d [11] followed by removal of UMI-identified PCR-duplicated using UMI-tools v0.5.5 [12]. Alignments with mapping qualities below 30 as well as short (< 40 base pairs) and secondary alignments were removed by SAMBAMBA v0.7.1 [13] prior to gene expression quantification using featureCounts v1.6.3 [14] and the ENSEMBL gene annotation (Homo_sapiens.GRCh38.92.gtf).

Gene quantification counts from two SARS-CoV-2 and two control samples with sufficient numbers of high quality read counts were processed by DESeq2 v1.28.1 [15] to calculate differential gene expression data. FDR-values were refined using the R-package fdrtool v1.2.15 [16]. Geneontology (GO) term analysis of significantly regulated genes (padj < 0.1) was performed using R-package clusterProfiler v3.16.0 [17].

Ranked gene lists for gene set enrichment analysis (GSEA) were generated using R-package fgsea v1.14.0 [18] and GSEA for KEGG pathway gene sets was performed using webgestalt [19] (<http://www.webgestalt.org>).

**Statistics:** Statistical analyses was performed by using Prism 8, version 8.4.1. (GraphPad Software, LLC)

**Supplementary Table S1:** Histopathological assessment of A. carotis communis samples performed on HE stained sections of formalin-fixed paraffin embedded tissue (FFPE).

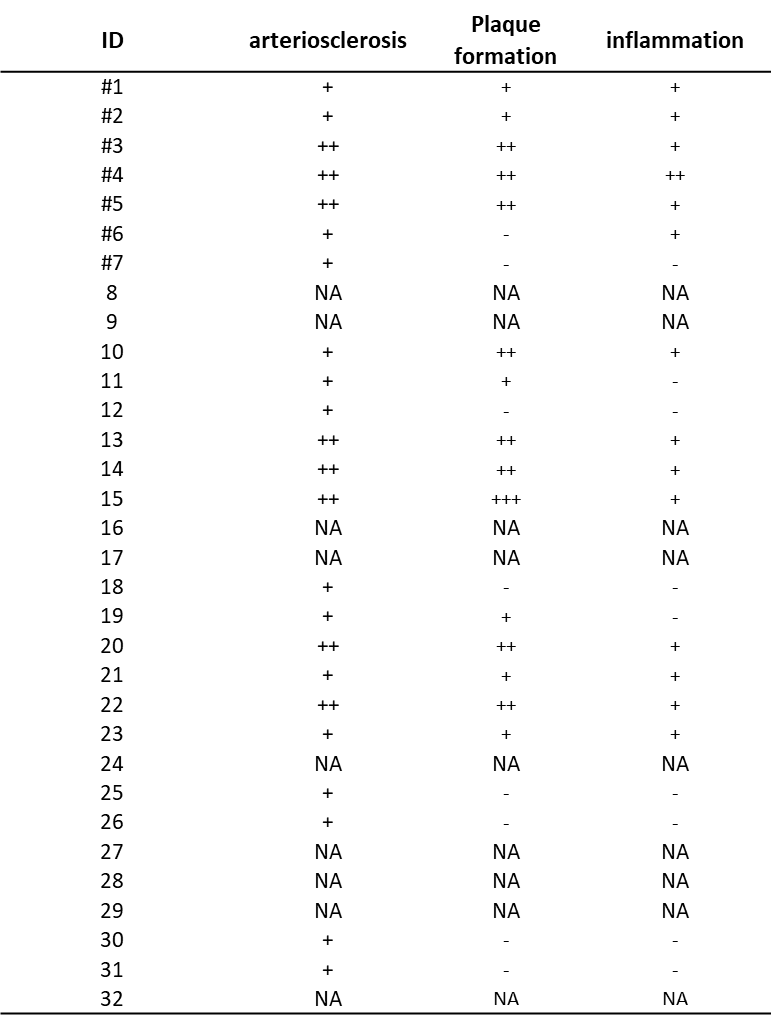

The degree of arteriosclerosis, plaque formation and vessel associated lymphocytic infiltration was assessed using a semi quantitative approach: none (-), slight (+), moderate (++), and severe (+++). Tissue samples from patients included in organ tropism and virus culture experiments are labelled with #.

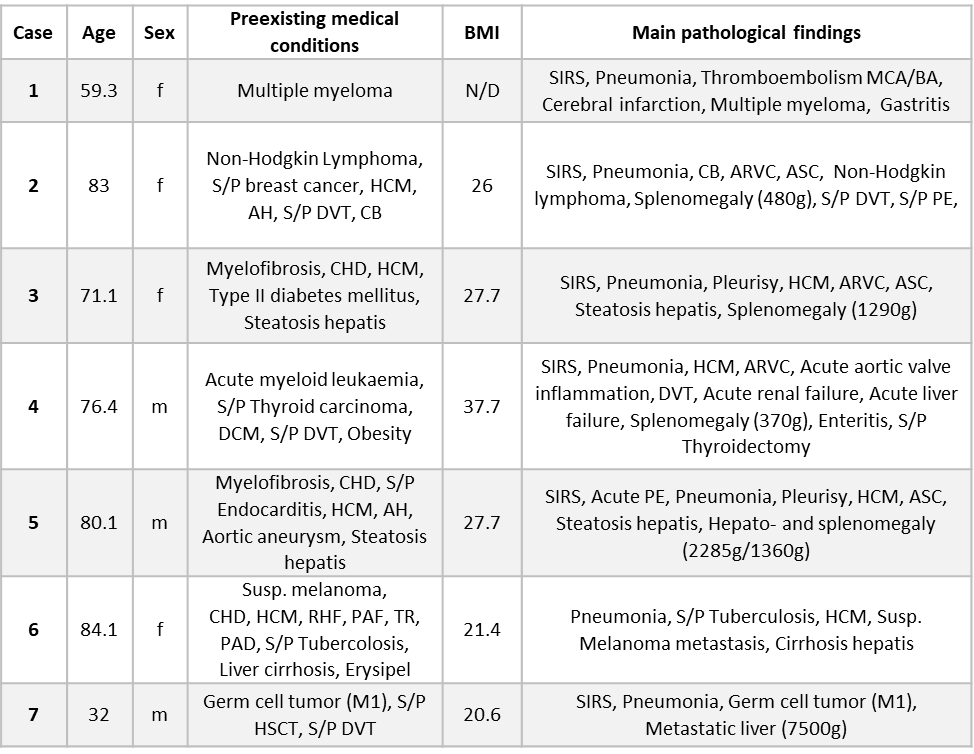
**Supplementary Table S2:** Summary of preexisting comorbidities and postmortem findings of 7 patients included for detailed analysis.

AH = arterial hypertension, ARVC = Arrhythmogenic right ventricular cardiomyopathy, ASC = Arteriosclerosis, BA = Basilar artery, CB = chronic bronchitis, CHD = Coronary heart disease, DCM = Dilatative cardiomyopathy, DVT = Deep venous thrombosis, HCM = Hypertrophic cardiomyopathy, HSCT = Hematopoietic stem cell transplantation, MCA = Middle cerebral artery, MODS = Multiorgan dysfunction syndrome, N/D= data not available; PAD = Peripheral artery disease, PAF = Paroxysmal atrial fibrillation, PE = Pulmonary embolism, RHF = Right heart failure, SIRS = Systemic inflammatory response syndrome, S/P = Status Post, Susp. = Suspected, TR =Tricuspid regurgitation.

**Supplementary Table S3:** Subgenomic RNA detection by targeted amplicon sequencing.

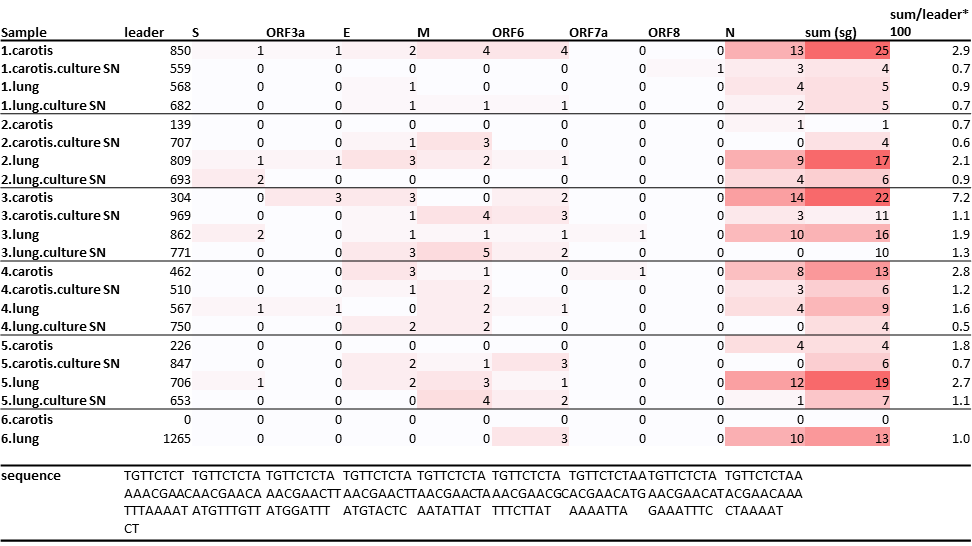

Shown are the number of amplicon reads spanning the leader sequence fused to the genome region encoding for structural and accessory proteins.

Sequences of the leader sequence of the individual are shown at the bottom. Sum sg represents total number of fragments subgenomic mRNAs across all samples.

**Supplementary Figure S1:** Clustering of viral variants of SARS-CoV-2 sequences recovered from tissue sections of A. carotis and lung tissue.

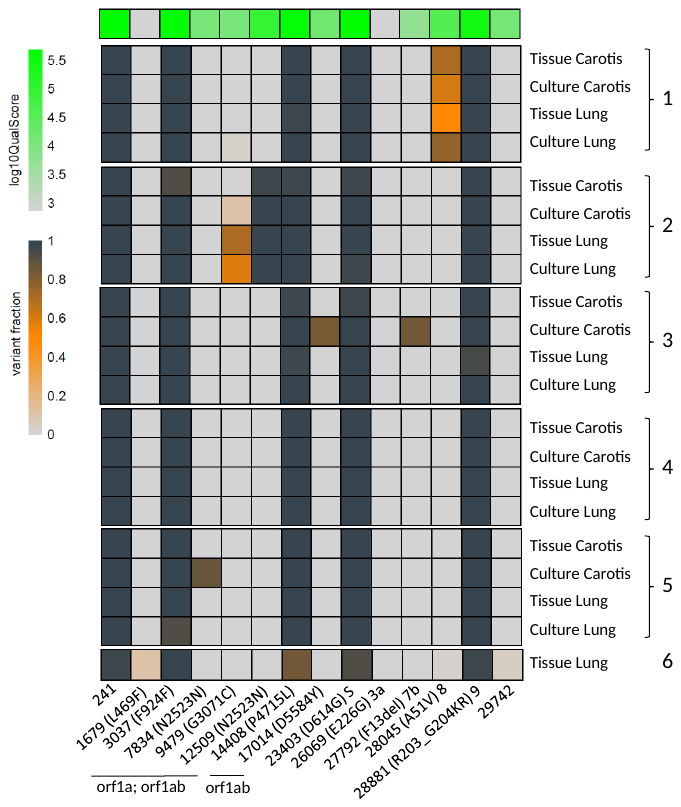

Viral sequences determined by amplicon sequencing were compared to sequences recovered from cell culture supernatant of Vero cells co-cultured with lysate of the diagnostic specimen. SARS-CoV-2 sequences from A. carotis and lung tissue from six autopsies, were compared to the reference sequence, NC_045521. Nucleotide positions of the observed variants are indicated at the bottom. Included are only variants with sufficient coverage (>10) and SNP being present in more than 33% of all reads in at least one sample. The frequency of variants is indicated by the heat map ranging from grey (reference), yellow to dark blue (variant). The quality score per individual site is indicated at the top.

**Supplementary Figure S2:** Control SARS-CoV-2 spike protein staining on tissue sections and infected Vero cells.

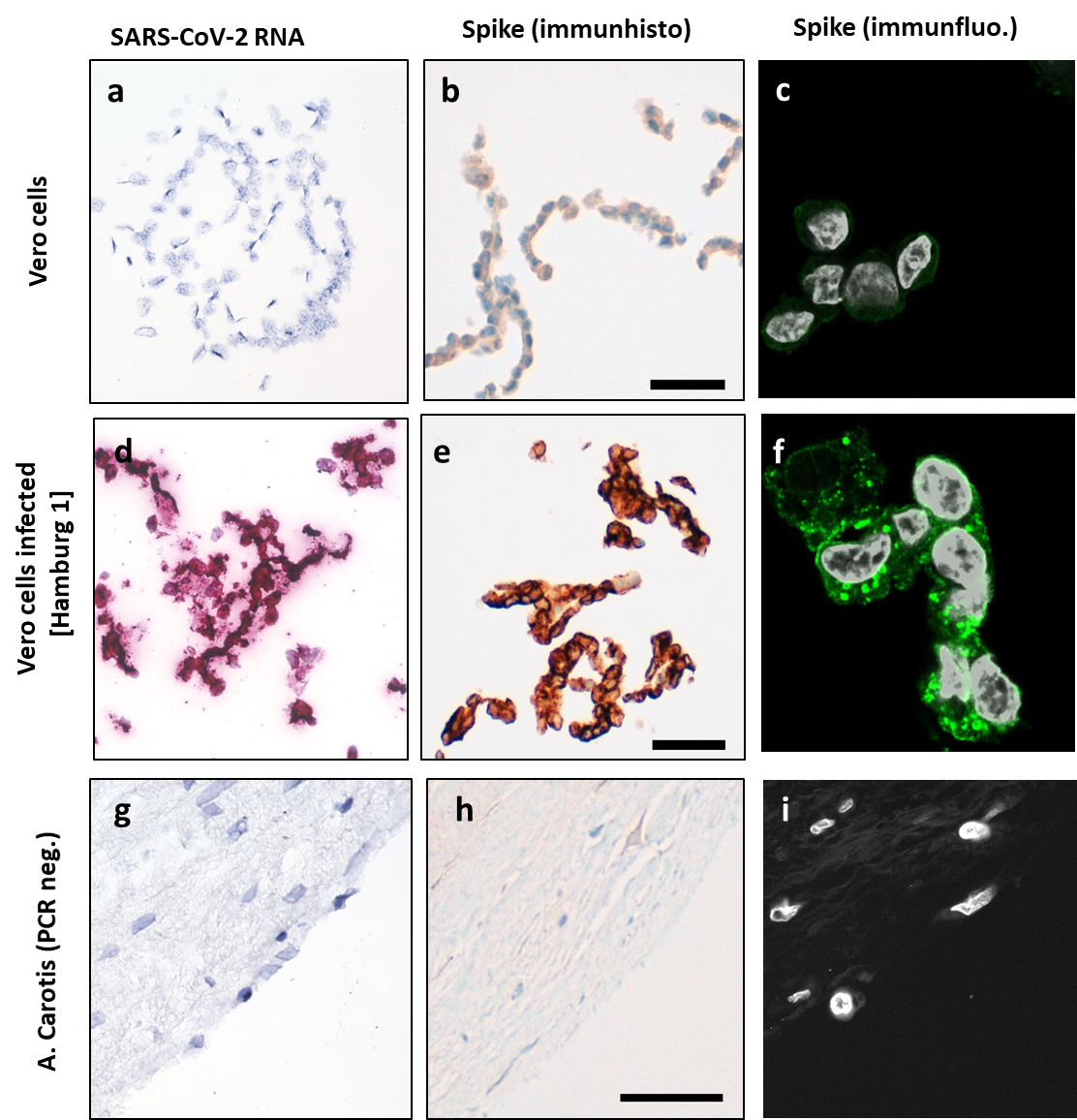

**Figure S2:** SARS-CoV-2 RNA in situ hybridization (pink; (a, d, g), spike protein staining by immunohistochemical staining (b, e, h) and indirect immunofluorescence (SARS-spike-protein in green, DNA in white), c, f, i of SARS-CoV-2 infected Vero cells (d, e, f). MOCK-infected Vero cells (a, b, c) and a SARS-CoV-2 PCR negative A. carotis sample (g, h, i) were used as negative controls. Scale bars are 25µm for a, b, d, e; 50µm for g, h and 10µm for c, f, and i.

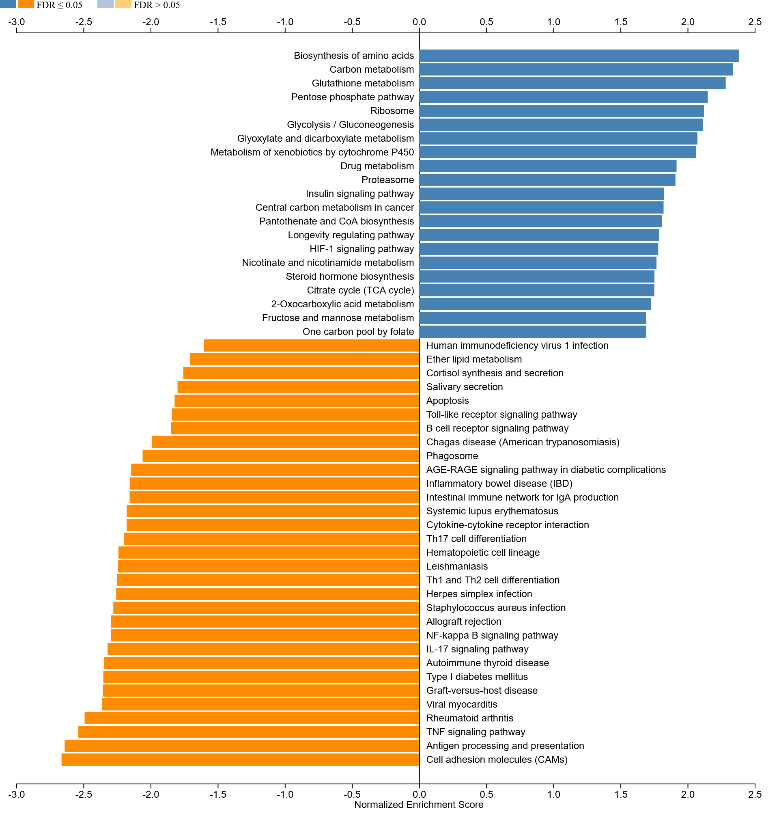
**Supplementary Figure S3:** Gene set enrichment analysis, GSEA.

Gene set enrichment analysis, GSEA, of KEGG PATHWAY categories.
